# Explicit modeling of ancestry improves polygenic risk scores and BLUP prediction

**DOI:** 10.1101/012005

**Authors:** Chia-Yen Chen, Jiali Han, David J. Hunter, Peter Kraft, Alkes L. Price

## Abstract

Polygenic prediction using genome-wide SNPs can provide high prediction accuracy for complex traits. Here, we investigate the question of how to account for genetic ancestry when conducting polygenic prediction. We show that the accuracy of polygenic prediction in structured populations may be partly due to genetic ancestry. However, we hypothesized that explicitly modeling ancestry could improve polygenic prediction accuracy. We analyzed three GWAS of hair color, tanning ability and basal cell carcinoma (BCC) in European Americans (sample size from 7,440 to 9,822) and considered two widely used polygenic prediction approaches: polygenic risk scores (PRS) and Best Linear Unbiased Prediction (BLUP). We compared polygenic prediction without correction for ancestry to polygenic prediction with ancestry as a separate component in the model. In 10-fold cross-validation using the PRS approach, the R^2^ for hair color increased by 66% (0.0456 to 0.0755; p<10^−16^), the R^2^ for tanning ability increased by % (0.0154 to 0.0344; p<10^−16^) and the liability-scale R^2^ for BCC increased by 68% (0.0138 to 0.0232; p<10^−16^) when explicitly modeling ancestry, which prevents ancestry effects from entering into each SNP effect and being over-weighted. Surprisingly, explicitly modeling ancestry produces a similar improvement when using the BLUP approach, which fits all SNPs simultaneously in a single variance component and causes ancestry to be under-weighted. We validate our findings via simulations, which show that the differences in prediction accuracy will increase in magnitude as sample sizes increase. In summary, our results show that explicitly modeling ancestry can be important in both PRS and BLUP prediction.

## Introduction

Genome-wide association studies (GWAS) have identified many single nucleotide polymorphisms (SNPs) associated with complex traits [Visscher et al. 2012]. Genetic prediction based on genome-wide significant SNPs from GWAS provides some predictive ability [Wray et al. 2007; Meigs et al. 2008; Kraft and Hunter 2009; So et al. 2011], but polygenic prediction, using genetic variants that do not achieve genome-wide significance, can improve prediction accuracy [de los Campos et al. 2010; Chatterjee et al. 2013; Dudbridge 2013]. Polygenic prediction has been applied to many complex traits, such as schizophrenia and height [Evans et al. 2009; Purcell et al. 2009; Wei et al. 2009; Bush et al. 2010; Lango Allen et al. 2010; Machiela et al. 2011; Makowsky et al. 2011; Stahl et al. 2012; Vazquez et al. 2012; Wei et al. 2013; Smoller et al. 2013; Rietveld et al. 2013; Ripke et al. 2014; Wood et al. 2014].

Polygenic prediction accuracy may be partly due to genetic ancestry, especially for complex traits that are known to be associated with ancestry. For example, differences in height between Northern Europeans and Southern Europeans (e.g. due to natural selection) explain 5% of the variance of height in European Americans from the Framingham Heart Study, accounting for most of the polygenic prediction accuracy in 1,997 unrelated individuals from this sample [Makowsky et al. 2011; Wray et al. 2013; Turchin et al. 2012]. Other traits that are well-known to be associated with ancestry in European-ancestry populations include hair color, eye color, skin pigmentation, skin cancer, multiple sclerosis, rheumatoid arthritis, type 1 diabetes, Crohn’s disease, Alzheimer’s disease, coagulation factor VII (FVII) plasma level, hemoglobin disorders, and educational attainment [Candille et al. 2012; Nan et al. 2009a; Rosati 2001; Cimmino et al. 1998; Patterson et al. 2009; Kenny et al. 2012; Panza et al. 2003; Bernardi et al. 1997; Angastiniotis and Modell 1998; Borjas 1994]. Thus, it is possible that polygenic prediction studies of these traits may likewise owe some of their accuracy to genetic ancestry [Evans et al. 2009; Wei et al. 2009; Bush et al. 2010; Makowsky et al. 2011; Stahl et al. 2012; Vazquez et al. 2012; Wei et al. 2013; Rietveld et al. 2013]. In genetic association studies, ancestry is a confounder that may cause false positive associations with no biological significance [Price et al. 2010]. On the other hand, if the ultimate goal is to conduct prediction in the same population, polygenic predictions that are partly due to ancestry may be useful. However, it is important to understand how much of polygenic prediction accuracy is due to ancestry, as this will affect prediction accuracy in other populations.

An open question is how to make use of ancestry information in polygenic prediction. Most studies of polygenic prediction ignore ancestry information, but this may not be optimal. We propose to correct for ancestry in the polygenic model using principal components, while incorporating associations between ancestry and the trait as a separate component in the prediction model. We hypothesize that explicit modeling of ancestry in this fashion can improve polygenic prediction accuracy, by preventing ancestry from entering into each SNP effect and being overweighted. In this study, we investigated the impact of explicit modeling of genetic ancestry on polygenic prediction accuracy by analyzing GWAS samples of European Americans for three pigmentation-related traits, including natural hair color (HC), childhood and adolescent tanning ability (TA), and basal cell carcinoma (BCC). We employed cross-validation schemes to ensure that improvements in prediction accuracy were not caused by over-fitting. We determined that explicitly modeling ancestry can significantly improve prediction accuracy across different polygenic prediction approaches, such as polygenic risk scores and best linear unbiased predictors. We note that although polygenic risk scores may not be the optimal prediction method when raw genotypes are available, we have chosen to include them as one of the main focuses of our analyses, because they are currently the most widely used approach for polygenic risk prediction in humans.

## Materials and Methods

### GWAS data

We performed analyses in 3 genome-wide association studies (GWAS) of hair color (HC), tanning ability (TA), and basal cell carcinoma (BCC). To obtain the GWAS samples, we pooled 7 GWAS on type 2 diabetes, coronary heart disease, kidney stone and breast cancer nested in the Nurses' Health Study (NHS) and the Health Professionals Follow-up Study (HPFS). The sample collection, genotyping and quality control for each individual GWAS were described in detail previously [Hunter et al. 2007; Nan et al. 2011; Qi et al. 2010; Rimm et al. 1991; Curhan and Taylor 2008; Taylor et al. 2005]. All samples are self-reported European Americans. These samples were genotyped with the Illumina HumanHap550 array, Illumina HumanHap610 Quad array, or Affymetrix 6.0 array. We included only SNPs genotyped in all 7 GWAS with call rate >99% and minor allele frequency >1% and excluded A/T and C/G SNPs to avoid strand ambiguity. The missing genotypes were mean-imputed in the subsequent analyses. The merged GWAS included 10,508 samples with 131,134 genotyped SNPs. From the merged GWAS, we created 3 GWAS for the target traits. Only samples with complete phenotype and self-reported ancestry information were included in each corresponding GWAS. We obtained 7,440 samples with HC, 9,822 samples with TA, and 2,086 BCC cases and 6,173 controls. All following analyses were repeated for each trait with the corresponding GWAS samples. We applied LD pruning by genomic position with a R^2^ threshold of 0.2 to obtain 71,557, 71,549 and 71,575 independent SNPs for HC, TA, and BCC, respectively.

In addition to the genotyped SNPs, we obtained imputed genotypes for variants that showed genome-wide significant association with HC, TA and BCC. This included 12 SNPs and one two-SNP haplotype associated with each of HC, TA and BCC and 6 additional SNPs associated with BCC only (Table S1) [Gerstenblith et al. 2010]. Note that some of these genome-wide significant SNPs were identified in the samples analyzed here, especially for TA; therefore, higher prediction accuracy based these genome-wide significant SNPs is expected [Han et al. 2008; Nan et al. 2009a; Nan et al. 2009b]. The imputation was done with using BEAGLE (for phasing) and MaCH using the 1000 Genomes Project Interim Phase I release as the reference panel [The 1000 Genomes Project Consortium 2012; Li et al. 2010; Browning and Browning 2007].

### Phenotype definitions

The phenotype information was collected from prospective questionnaires in NHS and HPFS. The natural hair color at 20 years of age was collected in 4 levels: blonde, light brown, dark brown, and black, ranging from 1 to 4. We excluded samples with red hair color, which is a distinct trait from other hair colors and is mainly determined by the *MC1R* gene [Han et al. 2008]. The childhood and adolescent tanning ability with more than 2 hours of sun light exposure was also collected in 4 levels: practically none, light tan, average tan, and deep tan, ranging from 1 to 4. A previous study showed that deeper tan denotes a better response to sun light exposure and is associated with lower risk of skin cancer [Nan et al. 2009a]. Both hair color and tanning ability were analyzed as quantitative traits. The BCC cases were identified through biennial self-report; the validation rate of BCC self-reports was 90% in validation studies in NHS and HPFS [Han et al. 2006; Hunter et al. 1990].

### Self-reported ancestry

We obtained self-reported ancestry groups from questionnaires. The original question on ancestry allows the participants to choose more than 1 category among all categories. We assigned participants into 1 of the 4 categories: Scandinavian, Southern European, Ashkenazi Jewish, and Other European (which includes samples with no self-reported European sub-population ancestry information). We calculated the mean and standard deviation for each trait by self-reported ancestry groups. We also tested for association between self-reported ancestry and each trait with a Wald test for HC and TA and a χ^2^ test and odds ratios with 95% confidence intervals for BCC.

### Inference of genetic ancestry

To infer genetic ancestry, we performed principal component analysis (PCA) with EIGENSTRAT software to obtain principal components as measure of genetic ancestry [Price et al. 2006; Patterson et al. 2006]. We performed PCA on the 3 GWAS and obtained the top 10 PCs specific to each GWAS. We plotted PC1 and PC2 to show the structure of these GWAS samples with color-coding by self-reported ancestry. To further investigate population structure, we projected the samples from Eastern European countries in the Population Reference Sample (POPRES) to the PC space derived from the TA GWAS samples, which contained most of our merged GWAS samples. The collections and methods for the Population Reference Sample (POPRES) are described by Nelson *et al.* (2008). The datasets used for the analyses described in this manuscript were obtained from dbGaP at http://www.ncbi.nlm.nih.gov/projects/gap/cgibin/study.cgistudy_id=phs000145.v1.p1 through dbGaP accession number phs000145.v1.p1. The PC projection was done by applying the SNP weights, which were calculated with principal components and eigenvalues derived from the TA GWAS samples, to the POPRES samples [Chen et al. 2013]. To compare the structure of POPRES samples and our GWAS samples, we plotted the projected PC1 and PC2 of POPRES samples with the PC1 and PC2 of the TA GWAS samples.

### Prediction using genetic and self-reported ancestry

We demonstrated the prediction ability of ancestry by fitting regression models with self-reported ancestry, PC1 and PC2, the top 5 PCs, and the top 5 PCs plus self-reported ancestry in all samples. R^2^ was calculated for each model and a likelihood ratio test (LRT) was used to compare models. We used R^2^ on the observed scale for HC and TA and R^2^ on the liability scale for BCC, with prevalence parameter set to 0.3 [Lear and Smith 1997; Lee et al. 2012]. The R^2^ on liability scale is a linear transformation of R^2^ on the observed scale, where the case-control status is taken as a continuous variable. We used linear regression for HC and TA and logistic regression for BCC. To control for over-fitting, we also employed 10-fold cross-validation to calculate R^2^: we built models by estimating model coefficients with the training set which includes 90% of the total sample size and applied the estimated model coefficients to the validation set, which includes 10% of the total sample size excluded from the training set, and calculated predicted phenotypes for the validation samples. We repeated this process 10 times and pooled the validation samples with predicted phenotypes to calculate out-of-sample R^2^ between predicted and true phenotypes.

### Prediction using Genetic Risk Scores (GRS) based on genome-wide significant SNPs

We built prediction models based on genetic risk scores (GRS) using genome-wide significant SNPs associated to HC, TA and BCC and compared GRS models to models based on both GRS and ancestry. In order to compare these models, we employed both 10-fold cross-validation and 10x9-fold nested cross-validation. We employed 10x9-fold nested cross-validation to calculate out-of sample R^2^, which accounts for model over-fitting, in addition to 10-fold cross-validation that only gives in-sample R^2^.

For 10-fold cross-validation, we estimated effect sizes of the known associated SNPs in the training sets by fitting the following models in the 90% training samples:

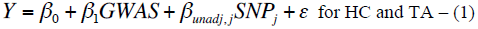

or

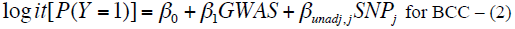

where SNP_j_ is the genotype of genome-wide significant SNP j and GWAS is a 6 × 1 vector of 6 indicator variables indicating which of the 7 individual GWAS the sample is from. We then applied the effect size estimates to the validation samples to construct GRS_unadj_. We calculated GRS_unadj_ for the 10% validation samples by

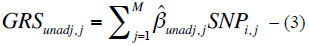

where 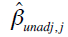 is the estimated regression coefficient using the training samples from model (1) and (2) and SNP_i,j_ is the genotype of validation sample i at SNP j. We also calculated GRS_adj_ by fitting models:

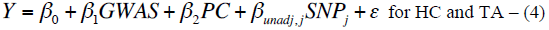

or

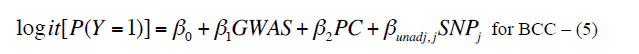

where SNP_j_ and GWAS are as previously defined and PC is a 5 × 1 vector of the top 5 PCs. The GRS_adj_ was then calculated as

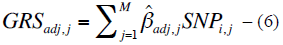

where 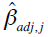 were estimated from model (4) and (5). We then pooled the samples from 10 mutually exclusive validation sets with GRS_unadj_ and GRS_adj_ constructed with estimates from their corresponding training samples and fitted models with GRS_unadj_, GRS_adj_, GRS_adj_ and the first 2 PCs, GRS_adj_ and the top 5 PCs, and GRS_adj_ and the top 5 PCs and self-reported ancestry. In-sample R^2^ and LRT p-values were calculated for these models.

For the 10x9-fold nested cross-validation, the GWAS samples were split into 2 parts with 90% of the total samples as the training set and 10% of the total samples as validation set in each fold of the nested cross-validation. For the 90% training set, we further split the samples into 2 training sets. The first training set had 80% of the total samples and the second training set had 10% of the total samples. We used the first training set to estimate effect sizes of the genome-wide significant SNPs and used those effect sizes to construct the GRS for the second training set as well as the validation set. We then fit prediction models that include GRS (and PCs/self-reported ancestry) in the second training set. We then applied the model coefficients from the second training set to the GRS (and PCs/self-reported ancestry) of the validation samples to obtain predicted phenotypes for the validation samples. For each of the 10 folds of validation samples, we repeated this model building for each of the 9 folds of first and second training sets and averaged the predicted phenotypes of validation samples across the 9 folds. With the pooled validation samples from the 10x9-fold nested cross-validation, we calculated out-of-sample R^2^ between true phenotypes and predicted phenotypes. The out-of-sample R^2^ were always calculated between a single predicted phenotype and the true phenotype of validation samples and the potential inflation of R^2^ due to increased number of predictors was eliminated.

### Polygenic prediction using Polygenic Risk Scores (PRS) based on genome-wide SNPs

For the polygenic model with explicit modeling of ancestry, we adjusted for the top 5 PCs when constructing the PRS and added the top 5 PCs to the polygenic model as additional covariates. We first did the analyses with 10-fold cross-validation using R^2^ and likelihood ratio test as figures of merit and then repeated the analyses in 10x9-fold nested cross-validation. Both 10-fold cross-validation and 10x9-fold nested cross-validation were performed in a similar fashion as described above for GRS model comparisons.

We adopted the polygenic risk score method to build polygenic prediction models [Evans et al. 2009; Purcell et al. 2009]. We used around 72,000 LD-pruned, independent genome-wide SNPs in PRS. To construct PRS without ancestry correction (PRS_unadj_), we fitted regression models (1) and (2) for each of the genome-wide SNPs individually using the 90% training samples. To construct PRS with ancestry correction (PRS_adj_), we fitted models (4) and (5) for each SNP individually. The 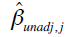 and 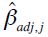 were used as weights to construct PRS_unadj_ and PRS_adj_,respectively. Models (3) and (6) were used to construct PRS, except that we included all genome-wide SNPs in the PRS. We also constructed PRS_unadj_ and PRS_adj_ using different numbers of genome-wide SNPs determined by the association p-value (calculated for each training set) for the SNP and the target phenotype: 5 × 10^−8^, 1 × 10^−7^, 1 × 10^−6^, 1 × 10^−5^, 1 × 10^−4^, 0.001, 0.01, 0.05, 0.1, 0.5, and 1.0. For BCC, we did not construct PRS with p-value threshold smaller than 1 × 10^−5^ because no SNP showed association p-value smaller than 1 × 10^−5^ for BCC in our data. We compared models with PRS_unadj_, PRS_adj_, PRS_adj_ and the top 5 PCs, and PRS_adj_ and the top 5 PCs and self-reported ancestry.

We further extended the polygenic models above to include both GRS_unadj_ and PRS_unadj,-gs_ or both GRS_adj_ and PRS_adj,-gs_. We excluded SNPs that showed genome-wide significant association with the target trait in previous GWAS (which are included in GRS), and SNPs in LD with those SNPs (pairwise R^2^ > 0.2), from PRS. The GRS and PRS were combined in the model by either summing the GRS and PRS to make one score or including GRS and PRS as two separate components of the model. We used in-sample R^2^ and LRT to compare these models in the 10-fold cross-validation and out-of-sample R^2^ in the 10x9-fold nested cross-validation.

### Polygenic prediction using Best Linear Unbiased Predictors (BLUP) based on genome-wide SNPs

In addition to PRS, we performed polygenic prediction based on BLUP by fitting linear mixed models to estimate genome-wide SNP effect sizes simultaneously [Yang et al. 2010]. We adopted 10-fold cross-validation to fit linear mixed models in the 90% training samples and output the random effects and fixed effects estimates with GCTA. The linear mixed model we used to obtain effect size estimates takes the general form:

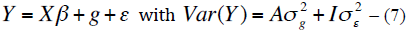

where *β* is a vector of fixed effects, g is the total genetic effects of the training samples with 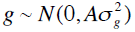 is the genetic relationship matrix derived from SNP genotypes, ε is the residual with 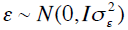 [Yang et al.2010]. In our case, the fixed effects included individual GWAS study indicator, PCs, and/or genome-wide significant SNPs. The effect sizes of SNPs included in PRS were calculated based on predicted g and training genotypes. We then apply the effect estimates to the 10% validation samples to construct PRS, GRS, and PCs. For BLUP_unadj_, BLUP_adj_, BLUP_unadj,-gs_ and BLUP_adj,-gs,_ the SNP effects were estimated based on BLUP. The GRS_unadj_, GRS_adj_ and PCs were entered the linear mixed models as fixed effects. The calculation of PRS in the validation samples is the same as models (3) and (6).

### Simulations comparing polygenic prediction using polygenic risk scores

It is widely known that polygenic prediction accuracy is predominantly determined by the training sample size. In our analysis using real GWAS samples, the training sample size is fixed. To examine the performance of different PRS approaches under different training sample sizes, we performed the following simulation study. We simulated a quantitative trait with 100,000 independent SNPs, where all SNPs were causal and the proportion of phenotypic variance explained by these SNPs was 0.5. We simulated sets of 1,000, 5,000, 10,000, 50,000, 100,000, 500,000, 1,000,000 training samples and 10,000 validation samples. Training samples and validation samples were each drawn from two subpopulations POP1 and POP2, with minor allele frequency equal to 0.50 in POP1 and drawn from a normal distribution with mean 0.50 and standard deviation 0.05 in POP2, so that the fixation index F_ST_(POP1,POP2)=0.005. The signs of marker effect sizes were flipped so that with probability 0.51, the allele with higher frequency in POP2 had positive effect, with the result that POP2 had systematically higher trait mean and subpopulation ancestry had an R^2^ of 0.044 with the trait. We compared three approaches: PRS_unadj_, PRS_adj_, and PRS_adj_ + ancestry, which is equivalent to PRS_adj_ + top PC except that ancestry is assumed to be inferred with perfect accuracy. (We did not consider additional PCs because the data was simulated to contain only one dimension of population structure.) Results were averaged across 100 simulations. We compared the simulation results to the results from our PRS analysis using real GWAS samples.

## Results

### Inference of genetic ancestry

We performed analyses in three genome-wide association studies (GWAS) of hair color (HC), tanning ability (TA), and basal cell carcinoma (BCC) with samples from the Nurses' Health Study (NHS) and the Health Professionals Follow-up Study (HPFS). A total of 7,440 samples with HC, 9,822 samples with TA, and 2,086 BCC cases and 6,173 controls were included, with around 72,000 SNPs after LD pruning in each GWAS. All samples are self-reported European Americans. We investigated the population structure in our GWAS samples using principal component analysis (PCA) [Price et al. 2006; Patterson et al. 2006], and compared the top principal components (PCs) to self-reported European ancestry. We first plotted PC1 and PC2 to investigate the population structure in the three GWAS (Figure 1). The PCA plot was color-coded with self-reported European ancestry, which has four categories: Scandinavian, Southern European, Ashkenazi Jewish, and Other European. We found that samples of self-reported Southern European ancestry had relatively lower PC1 values than samples of self-reported Scandinavian ancestry, while self-reported Ashkenazi Jewish samples had the lowest PC1 on average. In addition, the average PC1 and PC2 of Other European samples are close to the average PC1 and PC2 of Scandinavian samples, suggesting that Other European samples are predominantly of Northern European ancestry. The structure we identified in our GWAS sample is different from the canonical PC1 and PC2 in European American samples shown in Price *et al.* (2008) perhaps because the current data set has more samples of Eastern European ancestry than the samples analyzed in that paper. By projecting POPRES samples onto PC1 and PC2, we determined that samples with high PC2 value represent Eastern European ancestry (Figure S1) [Nelson et al. 2008]. This result suggests that lower PCs may capture additional structure. In the prediction analyses below, we include either PC1 and PC2 or the top 5 PCs; including more than 5 PCs (up to 10 PCs) did not change any of the results.

**Figure 1.**
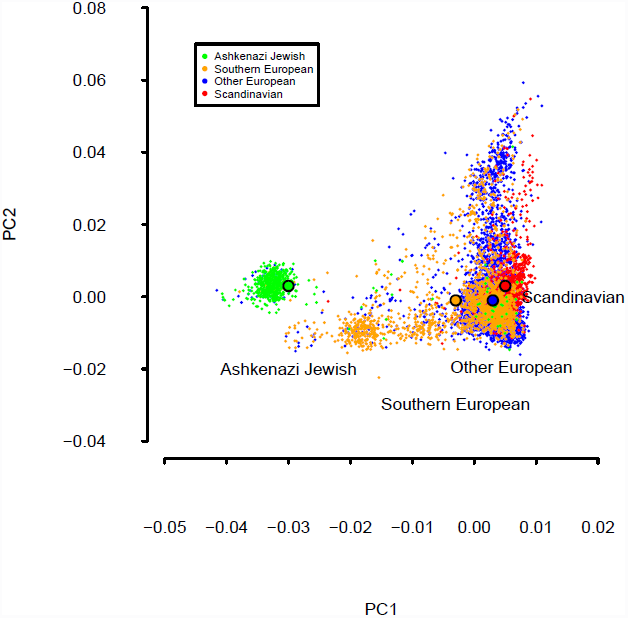
PCA plots for tanning ability GWAS samples, color-coded with self-reported ancestry. Black circles with color-coded interior denote average PC values for each self-reported ancestry. The tanning ability GWAS had the largest samples size among the 3 GWAS analyzed. Results for hair color and basal cell carcinoma GWAS samples, which are largely overlapped with the tanning ability GWAS samples, were extremely similar.

### Prediction using genetic and self-reported ancestry

Pigmentation-related traits are associated with ancestry, due to positive selection on lighter pigmentation in populations in regions of higher latitude [Sulem et al. 2007]. Indeed, our data showed that HC, TA and BCC are strongly associated with genome-wide ancestry and self-reported ancestry (Table 1). Compared with self-reported Scandinavian samples, Southern European samples on average have darker hair color (2.60 in Southern European vs. 2.08 in Scandinavian), increased tanning ability (2.67 in Southern European vs. 2.36 in Scandinavian), which corresponds to better protection against ultraviolet radiation, and lower risk of BCC (odds ratio [OR] for Scandinavian vs. Southern European (ref) = 1.075, 95% confidence interval [CI]: 1.036, 1.116). Ashkenazi Jewish samples on average had the darkest hair color (2.76), which is consistent with their lower PC1 value vs. South Europeans, but also had the lowest tanning ability (2.30) and highest risk of BCC among 4 self-reported ancestry groups (OR = 1.123, 95% CI: 1.074, 1.174), which are not consistent with their lower PC1 value vs. South Europeans. The Scandinavian samples had similar but slightly more extreme phenotypes than the Other European samples, consistent with the similar but slightly higher average PC1 for Scandinavian vs. Other European samples.

**Table 1.** Average PCs and phenotype values for self-reported ancestry groups in GWASs on hair color, tanning ability, and basal cell carcinoma

| Self-reported European ancestry | Average PCs (SD)<sup>a</sup> |  | Hair color |  | Tanning ability |  | Basal cell carcinoma |  |  |
| --- | --- | --- | --- | --- | --- | --- | --- | --- | --- |
|  | 1 | 2 | N | Average | N | Average | Cases | Control | OR (95% CI) |
| Ashkenazi Jewish | -0.030<br>(0.009) | 0.003<br>(0.004) | 425 | 2.76<br>(0.73) | 579 | 2.30<br>(0.87) | 338 | 155 | 1.123<br>(1.074, 1.174) |
| Southern European | -0.003<br>(0.010) | -0.001<br>(0.011) | 1325 | 2.60<br>(0.74) | 1700 | 2.67<br>(0.90) | 1151 | 285 | Reference |
| Other European<sup>b</sup> | 0.003<br>(0.005) | -0.001<br>(0.010) | 4963 | 2.39<br>(0.75) | 6559 | 2.54<br>(0.91) | 4074 | 1419 | 1.062<br>(1.035, 1.089) |
| Scandinavian | 0.005<br>(0.003) | 0.003<br>(0.009) | 727 | 2.08<br>(0.80) | 984 | 2.36<br>(0.91) | 610 | 227 | 1.075<br>(1.036, 1.116) |
| Total |  |  | **7440** |  | **9822** |  | **6173** | **2086** |  |

<sup>a</sup>Average PCs are derived from tanning ability GWAS samples, which has the largest sample size among the 3 GWAS analyzed here. Samples of the 3 GWAS are largely overlapped.
<sup>b</sup>Other European refers to European American samples without self-reported European sub-population ancestry information.

Motivated by these observations, we fitted prediction models based on ancestry. We first examined models based on self-reported ancestry only, where the in-sample R^2^ was 0.0397 for HC, 0.0081 for TA, and 0.0072 for BCC (Table 2). We then examined prediction models based on PCs only. The model based on top 5 PCs achieved R^2^ of 0.0724 for HC, 0.0350 for TA, and 0.0202 for BCC in the in-sample analysis (Table 2). We also showed that adding PC3 to PC5 in addition to PC1 and PC2 improved model prediction accuracy significantly for all 3 phenotypes (p < 10^–16^ for HC and TA; 1.93 × 10^−15^ for BCC). This suggests that including the subtle structure captured by later PCs provided additional value in predicting these phenotypes. However, including more than 5 PCs did not change any of the results. The R^2^ obtained by using self-reported ancestry are lower than the R^2^ obtained by using the top 5 PCs. Interestingly, we found including self-reported ancestry in the models in addition to the top 5 PCs further improved prediction accuracy (Table 2). The improvement was significant for HC (p = 1.32 × 10^−9^) and TA (p = 5.70 × 10^−3^) but not for BCC (p = 0.18) in the in-sample analysis. In addition, we showed that the added information in self-reported ancestry for HC (where this effect is strongest) is primarily due to self-reported Scandinavian ancestry, which is not completely captured by the Northwest-Southeast European cline of PC1 (Table S2). We also performed out-of-sample analyses using 10-fold cross-validation and obtained similar results (Table S3). Overall, we showed that both PCs and self-reported ancestry explained part of the phenotypic variation and can be used to build prediction models for these pigmentation-related traits. Furthermore, while models based on PCs have higher prediction accuracy than models based on self-reported ancestry, self-reported ancestry can still improve prediction accuracy over PCs.

**Table 2.**
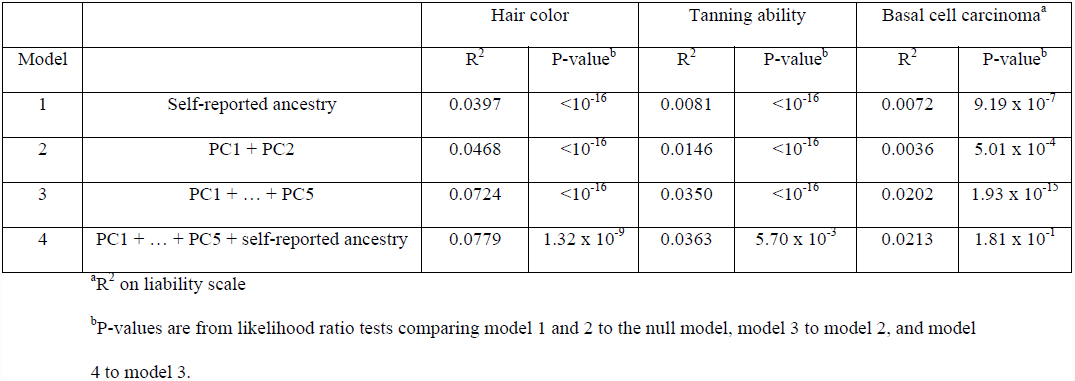
R^2^ and p-values from likelihood ratio test for prediction models using PCs and self-reported European ancestry in in-sample analysis using all samples

### Prediction using Genetic Risk Scores (GRS) based on genome-wide significant SNPs

To investigate the potential improvement of genetic risk prediction models by incorporating ancestry information, we compared genetic risk prediction models based on genetic risk scores (GRS) [Meigs et al. 2008]. We used imputed genotypes of SNPs shown to be genome-wide significant in previous GWAS to create GRS for each of the 3 phenotypes (Table S1). The models we compared included GRS without ancestry correction (GRS_unadj_), GRS with ancestry correction (adjusted for the top 5 PCs) (GRS_adj_), and GRS with explicit modeling of ancestry (GRS_adj_+PCs). In 10-fold cross-validation, GRSunadj attained an R^2^ of 0.2236 for HC (p < 10^-16^), 0.1311 for TA (p < 10^-16^), and 0.0266 (p < 10^-16^) for BCC (Table 3). The R^2^ of GRSadj were lower than the R^2^ of GRSunadj for HC and BCC, but slightly higher for TA (R^2^ = 0.1324). In 10x9-fold nested cross-validation, the R^2^ of GRS_adj_ were consistently lower than R^2^ of GRS_unadj_ for all 3 phenotypes (Table S3). The R^2^ of GRS_unadj_ and GRS_adj_ were much higher than the models based on ancestry. However, we showed that the prediction accuracy was improved (relative to either GRS_unadj_ or GRS_adj_) by including PCs in the GRSadj model (Table 3). We repeated the GRS_unadj_, GRS_adj_ and GRS_adj_+PCs model comparison by fitting all known genome-wide significant SNPs in one model simultaneously to obtain effect size estimates for constructing GRS (Table S4). The results showed similar patterns to the results from fitting genome-wide significant SNPs individually (Table 3). The improvement also remains for the out-of-sample R^2^ from 10x9-fold nested cross-validation (Table S3). We also calculated the R^2^ between the top 5 PCs and the predicted phenotypes based on GRS_unadj_ and GRS_adj_ (Table S5). We found that the R^2^ between PCs and GRSadj were only slightly lower than the R^2^ between PCs and GRSunadj, which suggests that the genome-wide significant SNPs used to calculate GRS have systematically different allele frequencies across European subpopulations, e.g. due to natural selection [Turchin et al. 2012]. In addition to PCs, we also showed that including self-reported ancestry further improved prediction accuracy on top of GRS_adj_ + PCs for HC (R^2^ from 0.2378 to 0.2423; p = 1.24 × 10^−9^) and TA (R^2^ from 0.1384 to 0.1397; p = 2.73 × 10^−3^), but not BCC (R^2^ from 0.0372 to 0.0385; p = 0.108) (Table S6).

**Table 3.**
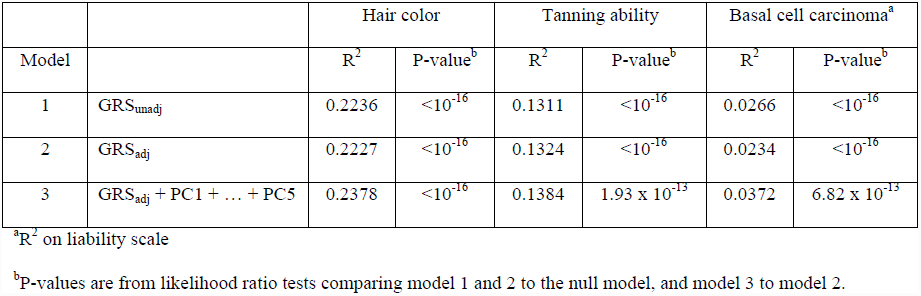
R^2^ and p-values from likelihood ratio test for prediction models using GWAS identified SNPs and PCs in 10-fold cross-validation

### Polygenic prediction using Polygenic Risk Scores (PRS) based on genome-wide SNPs

We investigated the impact of explicitly modeling ancestry on polygenic prediction by comparing 3 polygenic models: 1) the polygenic risk score (PRS) without ancestry correction (PRS_unadj_), 2) a polygenic risk score with correction for ancestry (PRS_adj_), and 3) a polygenic model with explicit modeling of ancestry, where we corrected for ancestry but added a separate ancestry component (PRS_adj_+PCs) [Purcell et al. 2009; Evans et al. 2009]. We compared PRS models with PRS that included all available SNPs across the genome after LD pruning. Results are displayed in Table 4. Comparing PRS_adj_+PCs to PRSunadj, the R^2^ improved 67% (from 0.0473 to 0.0789) for hair color, 114% (from 0.0162 to 0.0347) for tanning ability and 62% (from 0.0145 to 0.0235) for BCC. The model improvements were significant with p-value < 10^−16^. We found that PRSadj attained a lower R^2^ than the other models (0.0055 for HC; 0.0003 for TA; 0.0029 for BCC). Notably, the improvement in R^2^ of PRS_adj_+PCs over PRSadj was much larger than the R^2^ of PRS_unadj_ in each case, suggesting that PRS_unadj_ suffered from a mis-specified weighting of ancestry effects, which enter into each SNP effect in the unadjusted model. The low R^2^ of PRSadj is consistent with the relatively small sample size in our analyses [Chatterjee et al. 2013; Dudbridge 2013], and indicates that the prediction accuracy of PRSunadj was largely due to ancestry. Indeed, we observed R^2^ as large as 0.9250 between PC1 and PRS_unadj_ (Table S5). This is consistent with our simulations, in which the prediction R^2^ for PRS^unadj^ was consistently biased toward the same number, 0.044, which is the R^2^ between ancestry and the target trait. However, the prediction R^2^ of PRS_adj_ increased from 0.003 to 0.439 as the training sample size increased from 1,000 to 1,000,000, and PRS_adj_+ancestry consistently outperformed both PRS_unadj_ and PRS_adj_ (Table 5). Results for PRS_adj_ were close to the values predicted by theory in the absence of ancestry effects (see Equation (1) of Wray et al. 2013 Nat Rev Genet) [Wray et al. 2013]. These simulations show that explicit modeling of genetic ancestry can improve polygenic prediction accuracy, and that the magnitude of this improvement can become extremely large as sample sizes grow.

**Table 4.** R^2^ and p-values from likelihood ratio test for polygenic prediction using PRS, GRS and PCs in 10-fold cross-validation

(a) PRS included all SNPs available from genome-wide SNP chip after LD-pruning
| Model |  | Hair color |  | Tanning ability |  | Basal cell carcinoma<sup>a</sup> |  |
| --- | --- | --- | --- | --- | --- | --- | --- |
| | | $R^2$ | P-value<sup>b</sup> | $R^2$ | P-value<sup>b</sup> | $R^2$ | P-value<sup>b</sup> |
| 1 | PRS<sub>unadj</sub> | 0.0473 | $<10^{-16}$ | 0.0162 | $<10^{-16}$ | 0.0145 | $3.34 \times 10^{-15}$ |
| 2 | PRS<sub>adj</sub> | 0.0055 | $1.65 \times 10^{-10}$ | 0.0003 | $7.67 \times 10^{-2}$ | 0.0029 | $5.90 \times 10^{-4}$ |
| 3 | PRS<sub>adj</sub> + PC1 + ... + PC5 | 0.0789 | $<10^{-16}$ | 0.0347 | $<10^{-16}$ | 0.0235 | $<10^{-16}$ |
| 4 | PRS<sub>unadj</sub> + PC1 + ... + PC5 | 0.0745 | $<10^{-16}$ | 0.0344 | $<10^{-16}$ | 0.0226 | $1.36 \times 10^{-6}$ |

| Model |  | Hair color |  | Tanning ability |  | Basal cell carcinoma<sup>a</sup> |  |
| --- | --- | --- | --- | --- | --- | --- | --- |
| | | $R^2$ | P-value<sup>b</sup> | $R^2$ | P-value<sup>b</sup> | $R^2$ | P-value<sup>b</sup> |
| 5 | PRS<sub>unadj,-gs</sub> + GRS<sub>unadj</sub><sup>c</sup> | 0.2280 | $8.82 \times 10^{-11}$ | 0.1312 | 0.1776 | 0.0333 | $1.44 \times 10^{-8}$ |
| 6 | PRS<sub>adj,-gs</sub> + GRS<sub>adj</sub> | 0.2235 | $7.43 \times 10^{-3}$ | 0.1326 | 0.1820 | 0.0259 | $6.23 \times 10^{-4}$ |
| 7 | PRS<sub>adj,-gs</sub> + GRS<sub>adj</sub> + PC1 + ... + PC5 | 0.2388 | $<10^{-16}$ | 0.1386 | $2.45 \times 10^{-13}$ | 0.0397 | $5.46 \times 10^{-13}$ |
| 8 | PRS<sub>unadj,-gs</sub> + GRS<sub>unadj</sub> + PC1 + ... + PC5 | 0.2376 | $<10^{-16}$ | 0.1378 | $1.39 \times 10^{-14}$ | 0.0398 | $6.00 \times 10^{-6}$ |

<sup>a</sup> $R^2$ on liability scale
<sup>b</sup>P-values are from likelihood ratio tests comparing model 1 and 2 to the null model, model 3 to model 2, model 4 to model 1, model 5 and 6 to the corresponding models with only GRS, and model 7 to model 6, and model 8 to model 5.
<sup>c</sup>The $R^2$ for PRS<sub>unadj,-gs</sub> + GRS<sub>adj</sub> were 0.2284, 0.1327 and 0.0308 for hair color, tanning ability, basal cell carcinoma. -gs denotes excluding genome-wide significant SNPs.

**Table 5.**
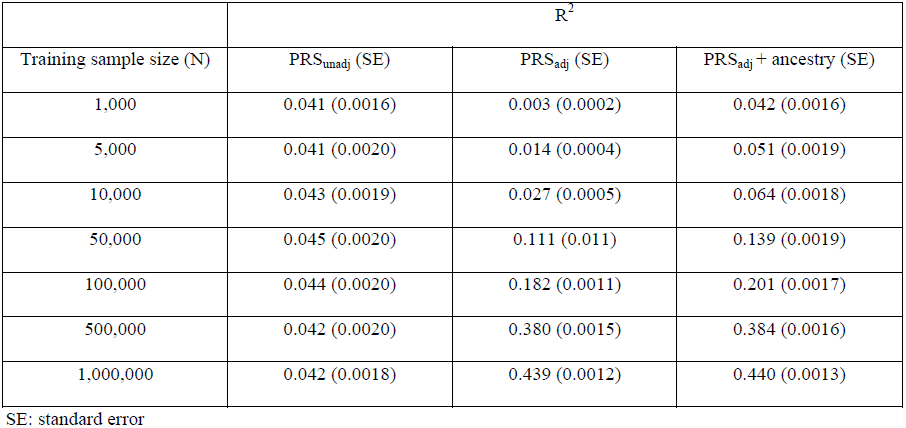
Comparison between prediction R^2^ of PRS_unadj_, PRS_adj_, and PRS_adj_ + ancestry in simulations with different training sample sizes The fluctuations in R^2^ of PRS_unadj_ are not statistically significant based on the number of simulations performed.

We also compared the 3 types of polygenic models with GRS_unadj_ or GRS_adj_ included as a component separate from PRS_unadj_ or PRS_adj_ in the model (PRS_unadj_ + GRS_unadj_, PRS_adj_ + GRS_adj_, and PRS_adj_ + GRS_adj_ + PCs). We excluded genome-wide significant SNPs (and SNPs in LD with known associated SNPs) from PRSunadj and PRS_adj_, since GRS based on genome-wide significant SNPs were included in the model as a separate component. Comparing the models with both PRS + GRS to the models with PRS only, the R^2^ in general improved and the improvements were consistent with the R^2^ from the GRS models (Table 4). The improvement in R^2^ due to explicit modeling of ancestry remained but with a smaller magnitude, similar to the analysis using GRS only (Table 3); similar relative results were obtained when using GRSadj in each model. For the PRS + GRS analyses, we also constructed GRS by fitting all known genome-wide significant SNPs in one model simultaneously to obtain effect size estimates (Table S7). The results showed similar patterns to the results from fitting genome-wide significant SNPs individually (Table 4). We also performed out-of-sample analyses using 10x9-fold nested cross-validation and obtained similar results (Table S3).

In addition to the 3 polygenic models described above, we evaluated the models of PRS_unadj_+PCs and PRS_unadj_+GRS_unadj_+PCs and found that these models attained lower R^2^ than PRS_adj_ + PCs and PRS_adj_+GRS_adj_+PCs (Table 4). This result may be caused by the fact that PRSunadj and PCs are correlated due to ancestry information in both, which is not optimally modeled. Finally, we considered models including self-reported ancestry in addition to PCs as ancestry measures. The polygenic models with explicit modeling of ancestry were further improved by self-reported ancestry for HC and TA, but not BCC (Table S6), consistent with predictions using ancestry only (Table 2).

**Figure 2.**
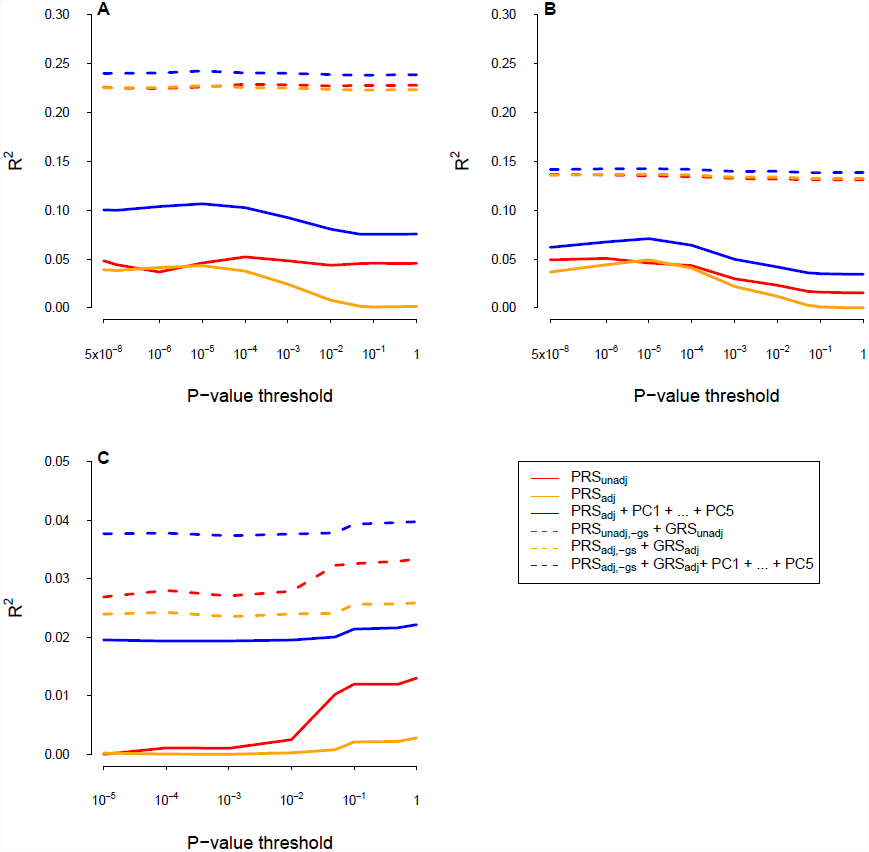
R^2^ for polygenic prediction using PRS with different association p-value thresholds, GRS and PCs in 10-fold cross-validation. (A): Hair color (B): Tanning ability (C): Basal cell carcinoma

In addition to using all independent SNPs across the genome to construct PRS, we applied a p-value threshold based on association tests between individual SNPs and corresponding phenotypes to select SNPs entering PRS. We used p-value thresholds from 5 × 10^−8^ to 1 and constructed PRS corresponding to each p-value threshold [Evans et al. 2009; Purcell et al. 2009]. We fitted the 3 types of polygenic models with PRS only and the 3 types of polygenic models with both PRS and GRS, using PRS subject to different p-value thresholds (Figure 2). The prediction accuracy for the polygenic models varied with p-value thresholds and showed different patterns for the 3 traits. The R^2^ of PRS_adj_+PCs model for HC and TA increased with more stringent p-value threshold, while R^2^ of PRSadj+PCs model for BCC remained stable. Overall, polygenic models with explicit modeling of ancestry had higher R^2^ than models without ancestry correction across all p-value thresholds applied to PRS. In addition, the difference between R^2^ from the model with explicit modeling of ancestry and the model without ancestry correction remained constant across p-value thresholds. For the PRS only models, the R^2^ changed with the p-value threshold for PRS but the magnitude of change was relatively small. For the models with both PRS and GRS components, the R^2^ were still improved with a smaller magnitude by explicitly modeling ancestry, compared with PRS only models. We also repeated the analyses in 10x9-fold nested cross-validation and the results were similar to 10-fold simple cross-validation (Figure S2). Overall, we showed that explicit modeling of genetic ancestry improves polygenic prediction accuracy, and that this pattern persists when including known associated SNPs via a GRS, and when using p-value thresholds.

### Polygenic prediction using Best Linear Unbiased Predictors (BLUP) based on genome-wide SNPs

As a comparison to PRS, which use SNP effect sizes obtained by fitting models for each SNP individually, we compared the 3 types of polygenic models based on best linear unbiased predictors (BLUP), which use SNP effect sizes obtained by fitting all SNPs using a linear mixed model [Henderson 1975; Haile-Mariam et al. 2013; de los Campos et al. 2013; Habier et al. 2013; Bolormaa et al. 2013]. We created 3 polygenic models: BLUP without ancestry correction (BLUP_unadj_); BLUP with ancestry correction (BLUP_adj_); and BLUP with explicit modeling of ancestry (BLUP_adj_+PCs); in BLUP_adj_ and BLUP_adj_+PCs, ancestry was modeled using fixed effects (see Methods). In general, the R^2^ of BLUP exhibited similar patterns as the R^2^ of PRS (Table 6). In particular, BLUP_adj_+PCs substantially outperformed BLUP_unadj_. As with PRS, the improvement in R^2^ of BLUP_adj_+PCs over BLUPadj was much larger than the R^2^ of BLUP_unadj_ in each case, suggesting that BLUP_unadj_ suffered from a mis-specified weighting of ancestry effects; although BLUP fits all SNPs simultaneously, it does not allow for different weights for different ancestry effects (e.g. PCs). BLUP_adj_ had low R^2^ due to small sample size, indicating that the prediction accuracy of BLUP_unadj_ was largely due to ancestry (Table S5). In models with BLUP and GRS as 2 separate components (where genome-wide significant SNPs were included in the GRS but not in BLUP), BLUP_adj_+GRS_adj_+PCs had significantly higher R^2^ than BLUP_unadj_+GRS_unadj_ (Table 6). Adding self-reported ancestry in the BLUP_adj_+GRS_adj_+PCs and the BLUP_adj_+PCs models improves the prediction accuracy significantly for HC and TA, but not BCC (Table S6).

**Table 6.** R^2^ and p-values from likelihood ratio test for polygenic prediction using BLUP, GRS and PCs in 10-fold cross-validation

(a) BLUP included all SNPs available from genome-wide SNP chip after LD-pruning
| Model |  | Hair color |  | Tanning ability |  | Basal cell carcinoma<sup>a</sup> |  |
| --- | --- | --- | --- | --- | --- | --- | --- |
| | | $R^2$ | P-value<sup>b</sup> | $R^2$ | P-value<sup>b</sup> | $R^2$ | P-value<sup>b</sup> |
| 1 | BLUP<sub>unadj</sub> | 0.0588 | $<10^{-16}$ | 0.0131 | $<10^{-16}$ | 0.0125 | $2.93 \times 10^{-13}$ |
| 2 | BLUP<sub>adj</sub> | 0.0045 | $5.86 \times 10^{-9}$ | 0.0001 | 0.385 | 0.0013 | 0.0112 |
| 3 | BLUP<sub>adj</sub> + PC1 + ... + PC5 | 0.0781 | $<10^{-16}$ | 0.0345 | $<10^{-16}$ | 0.0218 | $<10^{-16}$ |

| Model |  | Hair color |  | Tanning ability |  | Basal cell carcinoma<sup>a</sup> |  |
| --- | --- | --- | --- | --- | --- | --- | --- |
| | | $R^2$ | P-value<sup>b</sup> | $R^2$ | P-value<sup>b</sup> | $R^2$ | P-value<sup>b</sup> |
| 4 | BLUP<sub>unadj,-gs</sub> + GRS<sub>unadj</sub> | 0.2350 | $<10^{-16}$ | 0.1221 | $8.07 \times 10^{-1}$ | 0.0275 | $6.49 \times 10^{-7}$ |
| 5 | BLUP<sub>adj,-gs</sub> + GRS<sub>adj</sub> | 0.2258 | $8.10 \times 10^{-3}$ | 0.1233 | $2.97 \times 10^{-6}$ | 0.0208 | $5.52 \times 10^{-2}$ |
| 6 | BLUP<sub>adj,-gs</sub> + GRS<sub>adj</sub> + PC1 + ... + PC5 | 0.2413 | $<10^{-16}$ | 0.1308 | $<10^{-16}$ | 0.0359 | $3.83 \times 10^{-15}$ |

<sup>b</sup> $R^2$ on liability scale
<sup>b</sup>P-values are from likelihood ratio tests comparing model 1 and 2 to the null model, model 3 to model 2, model 4 and 5 to the corresponding models with only GRS, and model 6 to model 5.
<sub>-gs</sub> denotes excluding genome-wide significant SNPs.

## Discussion

Polygenic prediction using genome-wide SNPs has been widely applied to many complex traits. The most widely used polygenic prediction method is to construct a polygenic risk score (PRS), which is the sum of allele counts weighted by effect sizes estimated individually for each SNP, optionally restricting to SNPs with association p-value below a certain threshold [Chatterjee et al. 2013; Dudbridge 2013]. (It has previously been noted (Chatterjee *et al.* 2013) that other model selection and shrinkage methods, such as least absolute shrinkage and selection operator (LASSO)-type threshold methods that analyze all SNPs simultaneously, may modestly outperform the standard polygenic prediction approach). In previous studies using PRS, some of them adjusted for ancestry (PRS_adj_)[Purcell et al. 2009; Stahl et al. 2012; Smoller et al. 2013; Rietveld et al. 2013, PGC 2014 Nature], and some of them did not (PRS_unadj_) [Evans et al. 2009; Bush et al. 2010; Lango Allen et al. 2010; Machiela et al. 2011; Wood et al. 2014 Nat Genet]. However, a separate point is that modeling genetic ancestry as a separate component (PRS_adj_+PCs) can improve polygenic prediction accuracy relative to either PRS_unadj_ or PRS_adj_, as we showed for HC, TA and BCC; the same result is expected to hold for other traits associated to ancestry in subtly structured populations; this could explain why recent GWAS of height, which have reported that their association results may be impacted by very subtle population stratification [Lango Allen et al. 2010; Wood et al. 2014 Nat Genet], have attained prediction R2 using PRS that are lower than predicted by theory [Daetwyler et al. 2008 PLoS ONE]. In addition, the result still holds when including known associated SNPs via a genetic risk score (GRS), and when using p-value thresholds. In the European American data we analyzed, using 5 PCs to model ancestry improved our results relative to using 2 PCs. Surprisingly, including self-reported ancestry in addition to 5 PCs improved results further, as self-reported ancestry contains information about Scandinavian ancestry that is not well-captured by the PCs. Thus, PCs are not guaranteed to capture all ancestry information in subtly structured populations.

When applying methods that do not model genetic ancestry as a separate component (e.g. PRS_unadj_ or PRS_adj_), it is still of interest to understand how much of polygenic prediction accuracy is due to ancestry, as this will affect prediction accuracy in other populations. For example, Rietveld et al. reported a polygenic prediction R^2^ of 0.02 (p < 10^−16^) in an analysis of unrelated samples but a polygenic prediction R^2^ of only 0.003 (p = 0.001) in a within-family analysis immune to ancestry effects (see Table S25 of Rietveld et al.) [Rietveld et al. 2013]. Unmodeled population structure provides one plausible explanation for this discrepancy. We note that although Rietveld et al. adjusted for 4 PCs in their analyses (PRS_adj_), there is no guarantee that PCs capture all ancestry information in subtly structure populations. In addition, it is possible that PRS_adj_+PCs might attain a higher prediction R^2^ than PRS_adj_ in unrelated samples in the Rietveld et al. data, as we showed for HC, TA and BCC.

A previous study analyzed GWAS data from the Framingham Heart Study to compare prediction models for skin cancer risk using either pedigree information, or PCs, or polygenic prediction using Bayesian Lasso [Vazquez et al. 2012]. This study reported that polygenic prediction attained the highest prediction accuracy, PCs attained the next highest prediction accuracy, and pedigree information attained the lowest prediction accuracy. However this study did not explicitly model ancestry in the polygenic prediction model. Based on our results, it is possible that polygenic prediction with explicit modeling of ancestry might attain higher accuracy than the polygenic prediction approach employed by ref. [Vazquez et al. 2012]. It is also possible that combining pedigree information with polygenic predictions might improve prediction accuracy, although we have not explored that approach here[Haile-Mariam et al. 2013].

Alternatives to the widely used PRS approach for polygenic prediction include Best Linear Unbiased Prediction (BLUP) and its Bayesian extensions, which fit all markers simultaneously [de los Campos et al. 2013; Habier et al. 2013; Zhou et al. 2013; Bolormaa et al. 2013]. The potential benefits of explicitly modeling ancestry in this context may depend on the prior distribution of effect sizes that is assumed. Under an infinitesimal (Gaussian) prior, fitting all marker effects simultaneously may implicitly capture the effects of ancestry. However, our results showed that BLUP still showed an improvement in prediction accuracy when explicitly modeling ancestry. Although previous work has argued against including PCs as separate fixed effects in a mixed model when estimating components of heritability [Janss et al. 2012], and has shown that including PCs as separate fixed effects is generally not necessary when using standard mixed model methods for association [Yang et al. 2014], those studies did not consider prediction accuracy. Our recommendations for prediction are different from those recommendations for heritability estimation and association.

A limitation of our study is that the increased accuracy of PRS_adj_+PCs (relative to PRS_adj_) is contingent on validation samples having the same population structure as the training samples, as was the case in our cross-validation experiments. If validation samples do not have the same population structure then prediction accuracy will be lower, although PRS_adj_+PCs is still an appropriate strategy. (For example, if validation samples are from a homogeneous population, then PRS_adj_+PCs will produce predictions virtually identical to PRS_adj_.) Another potential limitation of our study is that it may not be feasible to compute PCs by running PCA on raw genotypes from the union of training and validation samples, as was done here; for example, this will not be feasible if only summary statistics are available for training samples. However, in this case, the top PCs for validation samples can be computed using weights derived from external reference panels (as projected PCs), as we have described previously [Chen et al. 2013]. We repeated the main analyses with top projected PCs derived for both training and validation samples using an external European American reference panel and obtained similar results to the results using PCs derived from raw genotypes of training and validation samples (Tables S8-S12). (We note that our main analyses use only training sample genotypes and phenotypes and validation sample genotypes to predict validation sample phenotypes, which is appropriate.)

We conclude by emphasizing the dependence of polygenic prediction accuracy on sample size. Both PRS_adj_ (with or without P-value thresholding to impose model selection) and BLUPadj attained low R2 in our analyses at modest sample sizes (<10,000), consistent with theoretical derivations [Daetwyler et al. 2008; Wray et al. 2013] and simulations suggesting that very large training sample sizes (100,000 or more) are needed to achieve high prediction accuracy [Chatterjee et al. 2013; Dudbridge 2013]. This limitation is in no way specific to the PRS approach. At larger sample sizes, it is likely that polygenic prediction will have significant clinical ramifications [Chatterjee et al. 2013; Dudbridge 2013]; indeed, prediction R2 has improved in recent very large studies [Ripke et al. 2014]. Notably, our simulations specifically show that the impact of explicit modeling of genetic ancestry will become extremely large as sample sizes grow.

## Acknowledgments

We are grateful to Mingfeng Zhang, Constance Chen, Naomi Wray and Peter Visscher for helpful discussions and to Arti Tandon for assistance with POPRES data. This work was supported by National Institutes of Health grant R01 HG006399 (C.Y.C. and A.L.P.).

## References

Angastiniotis M, Modell B. 1998. Global epidemiology of hemoglobin disorders. Ann N Y Acad Sci 850:251–69.

Bernardi F, Arcieri P, Bertina RM, Chiarotti F, Corral J, Pinotti M, Prydz H, Samama M, Sandset PM, Strom R and others. 1997. Contribution of Factor VII Genotype to Activated FVII Levels : Differences in Genotype Frequencies Between Northern and Southern European Populations. Arterioscler Thromb Vasc Biol 17:2548–553.

Bolormaa S, Pryce JE, Kemper K, Savin K, Hayes BJ, Barendse W, Zhang Y, Reich CM, Mason BA, Bunch RJ, and others. 2013. Accuracy of prediction of genomic breeding values for residual feed intake, carcass and meat quality traits in Bos taurus, Bos indicus and composite beef cattle. J Anim Sci 91:3088–104.

Borjas GJ. 1994. The Economics of Immigration. J Econ Lit 32:1667–717.

Browning SR, Browning BL, 2007. Rapid and accurate haplotype phasing and missing-data inference for whole-genome association studies by use of localized haplotype clustering. Am J Hum Genet 81:1084–97.

Bush WS, Sawcer SJ, Jager PLde, Oksenberg JR, McCauley JL, Pericak-Vance Ma, Haines JL. 2010. Evidence for polygenic susceptibility to multiple sclerosis--the shape of things to come. Am J Hum Genet 86:621–5.

Candille SI, Absher DM, Beleza S, Bauchet M, McEvoy B, Garrison Na, Li JZ, Myers RM, Barsh GS, Tang H and others. 2012. Genome-wide association studies of quantitatively measured skin, hair, and eye pigmentation in four European populations. PLoS One 7, 10, e48294.

Chatterjee N, Wheeler B, Sampson J, Hartge P, Chanock SJ, Park J-H. 2013. Projecting the performance of risk prediction based on polygenic analyses of genome-wide association studies. Nat Genet 45, 4, 400–5, 405e1–3.

Chen C-Y, Pollack S, Hunter DJ, Hirschhorn JN, Kraft P, Price AL. 2013. Improved ancestry inference using weights from external reference panels. Bioinformatics 29:1399–406.

Cimmino Ma, Parisi M, Moggiana G, Mela GS, Accardo S. 1998. Prevalence of rheumatoid arthritis in Italy: the Chiavari Study. Ann Rheum Dis 57:315–8.

Curhan GC, Taylor EN. 2008. 24-H Uric Acid Excretion and the Risk of Kidney Stones. Kidney Int 73:489–96.

Daetwyler HD, Villanueva B, Woolliams Ja, 2008. Accuracy of predicting the genetic risk of disease using a genome-wide approach. PLoS One 3:e3395.

Dudbridge F. 2013. Power and Predictive Accuracy of Polygenic Risk Scores. PLoS Genet 9, 3, e1003348.

Evans DM, Visscher PM, Wray NR. 2009. Harnessing the information contained within genome-wide association studies to improve individual prediction of complex disease risk. Hum Mol Genet 18:3525–31.

Gerstenblith MR, Shi J, Landi MT. 2010. Genome-wide association studies of pigmentation and skin cancer: a review and meta-analysis. Pigment Cell Melanoma Res 23:587–606.

Habier D, Fernando RL, Garrick DJ. 2013. Genomic-BLUP Decoded: A Look into the Black Box of Genomic Prediction. Genetics 194:597–607.

Haile-Mariam M, Nieuwhof GJ, Beard KT, Konstatinov KV, Hayes BJ. 2013. Comparison of heritabilities of dairy traits in Australian Holstein-Friesian cattle from genomic and pedigree data and implications for genomic evaluations. J Anim Breed Genet 130:20–31.

Han J, Kraft P, Colditz GA, Wong J, Hunter DJ. 2006. Melanocortin 1 receptor variants and skin cancer risk. Int J Cancer 119:1976–84.

Han J, Kraft P, Nan H, Guo Q, Chen C, Qureshi A, Hankinson SE, Hu FB, Duffy DL, Zhao ZZ and others. 2008. A genome-wide association study identifies novel alleles associated with hair color and skin pigmentation. PLoS Genet 4:e1000074.

Henderson CR. 1975. Best linear unbiased estimation and prediction under a selection model. Biometrics 31:423–47.

Hunter DJ, Colditz GA, Stampfer MJ, Rosner B, Willett WC, Speizer FE. 1990. Risk factors for basal cell carcinoma in a prospective cohort of women. Ann Epidemiol 1:13–23.

Hunter DJ, Kraft P, Jacobs KB, Cox DG, Yeager M, Hankinson SE, Wacholder S, Wang Z, Welch R, Hutchinson A and others. 2007. A genome-wide association study identifies alleles in FGFR2 associated with risk of sporadic postmenopausal breast cancer. Nat Genet 39:870–4.

Janss L, de Los Campos G, Sheehan N, Sorensen D. 2012. Inferences from genomic models in stratified populations. Genetics 192:693–704.

Kenny EE, Pe’er I, Karban A, Ozelius L, Mitchell AA, Ng SM, Erazo M, Ostrer H, Abraham C, Abreu MT and others. 2012. A genome-wide scan of Ashkenazi Jewish Crohn’s disease suggests novel susceptibility loci. PLoS Genet 8:e1002559.

Kraft P, Hunter DJ. 2009. Genetic risk prediction--are we there yet? N Engl J Med 360:1701–3.

Lango Allen H, Estrada K, Lettre G, Berndt SI, Weedon MN, Rivadeneira F, Willer CJ, Jackson AU, Vedantam S, Raychaudhuri S. 2010. Hundreds of variants clustered in genomic loci and biological pathways affect human height. Nature 467:832–8.

Lear JT, Smith AG. 1997. Basal cell carcinoma. Postgrad Med J 73:538–42.

Lee SH, Goddard ME, Wray NR, Visscher PM. 2012. A better coefficient of determination for genetic profile analysis. Genet Epidemiol 36:214–24.

Li Y, Willer CJ, Ding J, Scheet P, Abecasis GR. 2010. MaCH: using sequence and genotype data to estimate haplotypes and unobserved genotypes. Genet Epidemiol 34:816–34.

De los Campos G, Gianola D, Allison DB. 2010. Predicting genetic predisposition in humans: the promise of whole-genome markers. Nat Rev Genet 11:880–6.

De los Campos G, Vazquez AI, Fernando R, Klimentidis YC, Sorensen D. 2013. Prediction of Complex Human Traits Using the Genomic Best Linear Unbiased Predictor. PLoS Genet 9:e1003608.

Machiela MJ, Chen C-Y, Chen C, Chanock SJ, Hunter DJ, Kraft P. 2011. Evaluation of polygenic risk scores for predicting breast and prostate cancer risk. Genet Epidemiol 35:506–14.

Makowsky R, Pajewski NM, Klimentidis YC, Vazquez AI, Duarte CW, Allison DB, de los Campos G. 2011. Beyond missing heritability: prediction of complex traits. PLoS Genet 7:e1002051.

Meigs JB, Shrader P, Sullivan LM, McAteer JB, Fox CS, Dupuis J, Manning AK, Florez JC, Wilson PWF, D’Agostino RB and others. 2008. Genotype score in addition to common risk factors for prediction of type 2 diabetes. N Engl J Med 359:2208–19.

Nan H, Kraft P, Hunter DJ, Han J. 2009a. Genetic variants in pigmentation genes, pigmentary phenotypes, and risk of skin cancer in Caucasians. Int J Cancer 125:909–17.

Nan H, Kraft P, Qureshi AA, Guo Q, Chen C, Hankinson SE, Hu FB, Thomas G, Hoover RN, Chanock S. 2009b. Genome-wide association study of tanning phenotype in a population of European ancestry. J Invest Dermatol 129:2250–7.

Nan H, Xu M, Kraft P, Qureshi AA, Chen C, Guo Q, Hu FB, Curhan G, Amos CI, Wang L-E and others. 2011. Genome-wide association study identifies novel alleles associated with risk of cutaneous basal cell carcinoma and squamous cell carcinoma. Hum Mol Genet 20:3718–24.

Nelson MR, Bryc K, King KS, Indap A, Boyko AR, Novembre J, Briley LP, Maruyama Y, Waterworth DM, Vollenweider P. 2008. The Population Reference Sample, POPRES : A Resource for Population, Disease, and Pharmacological Genetics Research. Am J Hum Genet 83:347–58.

Panza F, Solfrizzi V, D’Introno A, Colacicco AM, Capurso C, Capurso A, Kehoe PG. 2003. Shifts in angiotensin I converting enzyme insertion allele frequency across Europe: implications for Alzheimer’s disease risk. J Neurol Neurosurg Psychiatry 74:1159–61.

Patterson CC, Dahlquist GG, Gyürüs E, Green A, Soltész G. 2009. Incidence trends for childhood type 1 diabetes in Europe during 1989-2003 and predicted new cases 2005-20: a multicentre prospective registration study. Lancet 373:2027–33.

Patterson N, Price AL, Reich D. 2006. Population structure and eigenanalysis. PLoS Genet 2:e190.

Price AL, Butler J, Patterson N, Capelli C, Pascali VL, Scarnicci F, Ruiz-Linares A, Groop L, Saetta A a, Korkolopoulou P and others. 2008. Discerning the Ancestry of European Americans in Genetic Association Studies. PLoS Genet 4:e236.

Price AL, Patterson NJ, Plenge RM, Weinblatt ME, Shadick Na, Reich D. 2006. Principal components analysis corrects for stratification in genome-wide association studies. Nat Genet 38:904–9.

Price AL, Zaitlen NA, Reich D, Patterson N. 2010. New approaches to population stratification in genome-wide association studies. Nat Rev Genet 11:459–63.

Purcell SM, Wray NR, Stone JL, Visscher PM, O’Donovan MC, Sullivan PF, Sklar P. 2009. Common polygenic variation contributes to risk of schizophrenia and bipolar disorder. Nature 460:748–52.

Qi L, Cornelis MC, Kraft P, Stanya KJ, Linda Kao WH, Pankow JS, Dupuis J, Florez JC, Fox CS, Paré G and others. 2010. Genetic variants at 2q24 are associated with susceptibility to type 2 diabetes. Hum Mol Genet 19:2706–15.

Rietveld CA, Medland SE, Derringer J, Yang J, Esko T, Martin NW, Westra H-J, Shakhbazov K, Abdellaoui A, Agrawal A and others. 2013. GWAS of 126,559 individuals identifies genetic variants associated with educational attainment. Science 340:1467–71.

Rimm EB, Giovannucci EL, Willett WC, Colditz Ga, Ascherio A, Rosner B, Stampfer MJ. 1991. Prospective study of alcohol consumption and risk of coronary disease in men. Lancet 338:464–8.

Ripke S, Neale BM, Corvin A, Walters JTR, Farh K-H, Holmans PA, Lee P, Bulik-Sullivan B, Collier DA, Huang H and others. 2014. Biological insights from 108 schizophrenia-associated genetic loci. Nature. 511:421–7.

Rosati G. 2001. The prevalence of multiple sclerosis in the world: an update. Neurol Sci 22:117–39.

Smoller JW, Craddock N, Kendler K, Lee PH, Neale BM, Nurnberger JI, Ripke S, Santangelo S, Sullivan PF. 2013. Identification of risk loci with shared effects on five major psychiatric disorders: a genome-wide analysis. Lancet 381:1371–9.

So H-., Kwan JSH, Cherny SS, Sham PC. 2011. Risk prediction of complex diseases from family history and known susceptibility loci, with applications for cancer screening. Am J Hum Genet 88:548–65.

Stahl EA, Wegmann D, Trynka G, Gutierrez-Achury J, Do R, Voight BF, Kraft P, Chen R, Kallberg HJ, Kurreeman FS and others. 2012. Bayesian inference analyses of the polygenic architecture of rheumatoid arthritis. Nat Genet 44:483–9.

Sulem P, Gudbjartsson DF, Stacey SN, Helgason A, Rafnar T, Magnusson KP, Manolescu A, Karason A, Palsson A, Thorleifsson G and others. 2007. Genetic determinants of hair, eye and skin pigmentation in Europeans. Nat Genet 39:1443–52.

Taylor E, Stampfer M, Curhan G. 2005. Obesity, weight gain, and the risk of kidney stones. J Am Med Assoc 293:455–62.

The 1000 Genomes Project Consortium. 2012. An integrated map of genetic variation from 1,092 human genomes. Nature 491:56–65.

Turchin MC, Chiang CWK, Palmer CD, Sankararaman S, Reich D, Hirschhorn JN. 2012. Evidence of widespread selection on standing variation in Europe at height-associated SNPs. Nat Genet 44:1015–9.

Vazquez AI, de Los Campos G, Klimentidis YC, Rosa GJM, Gianola D, Yi N, Allison DB. 2012. A comprehensive genetic approach for improving prediction of skin cancer risk in humans. Genetics 192:1493–502.

Visscher PM, Brown MA, McCarthy MI, Yang J. 2012. Five Years of GWAS Discovery. Am J Hum Genet 90:7–24.

Wei Z, Wang K, Qu H-Q, Zhang H, Bradfield J, Kim C, Frackleton E, Hou C, Glessner JT, Chiavacci R and others. 2009. From disease association to risk assessment: an optimistic view from genome-wide association studies on type 1 diabetes. PLoS Genet 5:e1000678.

Wei Z, Wang W, Bradfield J, Li J, Cardinale C, Frackelton E, Kim C, Mentch F, Steen KV, Visscher PM and others. 2013. Large sample size, wide variant spectrum, and advanced machine-learning technique boost risk prediction for inflammatory bowel disease. Am J Hum Genet 92:1008–12.

Wood AR, Esko T, Yang J, Vedantam S, Pers TH, Gustafsson S, Chu AY, Estrada K, Luan J, Kutalik Z and others. 2014. Defining the role of common variation in the genomic and biological architecture of adult human height. Nat Genet 46:1173–86.

Wray NR, Goddard ME, Visscher PM, 2007. Prediction of individual genetic risk to disease from genome-wide association studies. Genome Res 17:1520–8.

Wray NR, Yang J, Hayes BJ, Price AL, Goddard ME, Visscher PM. 2013. Pitfalls of predicting complex traits from SNPs. Nat Rev Genet 14:507–15.

Yang J, Lee SH, Goddard ME, Visscher PM. 2010. GCTA: A Tool for Genome-wide Complex Trait Analysis. Am J Hum Genet 88:76–82.

Yang J, Zaitlen N, Goddard ME, Visscher PM, Price AL. 2014. Advantages and pitfalls in the application of mixed-model association methods. Nat Genet 46:100–6.

Zhou X, Carbonetto P, Stephens M. 2013. Polygenic modeling with bayesian sparse linear mixed models. PLoS Genet 9:e1003264.

